# Thiophosphopeptides Instantly Targeting Golgi Apparatus and Selectively Killing Cancer Cells

**DOI:** 10.1101/2021.02.13.431079

**Authors:** Weiyi Tan, Qiuxin Zhang, Jiaqing Wang, Meihui Yi, Hongjian He, Bing Xu

## Abstract

Golgi apparatus is emerging as a key signaling hub of cells, but there are few approaches for targeting Golgi and selectively killing cancer cells. Here we show an unexpected result that changing an oxygen atom of the phosphoester bond in phospho-peptides by a sulfur atom enables instantly targeting Golgi apparatus (GA) and selectively killing cancer cells by enzymatic self-assembly. Specifically, conjugating cysteamine S-phosphate to the C-terminal of a self-assembling peptide generates a thiophosphopeptide. Being a substrate of alkaline phosphatase (ALP), the thiophosphopeptide undergoes rapid ALP-catalyzed dephosphorylation to form a thiopeptide that self-assembles. The thiophosphopeptide enters cells via caveolin-mediated endocytosis and macropinocytosis and instantly accumulates in GA because of dephosphorylation and formation of disulfide bonds in Golgi. Moreover, the thiophosphopeptide, targeting Golgi, potently and selectively inhibits cancer cells (e.g., HeLa) with the IC50 (about 3 μM), which is an order of magnitude more potent than that of the parent phosphopeptide. This work, as the first report of thiophosphopeptide for targeting Golgi, illustrates a new molecular platform for designing enzyme responsive molecules that target subcellular compartment for functions.

---

Golgi apparatus (GA), a stack of flattened membrane-enclosed disks that are dynamically regulated during cell cycles in mammalian cells, is considered as the “heart” of intracellular transportation.^1-2^ Increasing numbers of studies have revealed that Golgi is a hub for different signaling pathways that drive the survival and migration of cancer cells.^3-5^ Although Golgi is emerging as an important target for cancer therapy, there are, however, few approaches for targeting Golgi.^6-7^ While Golgi mannosidase II inhibitors are able to inhibit cancer cells, the selectivity^8^ or efficacy^6^ of the inhibitors remains to be improved. In addition, several studies reported the imaging of Golgi, including the commercial dyes for staining Golgi,^9^ a smart “off−on” fluorescence probe for imaging the Golgi in cancer cells,^10^ and carbon quantum dots localizing at Golgi.^11^ These imaging agents, however, require 30 minutes^10^ or longer incubation time^10^ or pretreatment,^9^ and they have yet to lead to the approach for selectively inhibiting the cancer cells. Thus, there is an unmet need of targeting Golgi to inhibit cancer cells.

During our study of enzymatic noncovalent synthesis (ENS),^12^ we changed an oxygen atom of the phosphoester bond in a phosphopeptide (**pO1**) by a sulfur atom to make **pS1** for fast enzymatic self-assembly. Our studies show that **pS1** undergoes rapid dephosphorylation catalyzed by ALP to form **S1** that assembles. Unexpectedly, treating HeLa cells with **pS1** shows that **S1** instantly accumulates at Golgi of the HeLa cells at the concentration as low as 500 nM. Such an enzymatic accumulation of Golgi (Scheme 1) is proportional to both the concentration of **pS1** and the time of incubation. Similar rapid enzymatic accumulation also takes places in the Golgi of several other cells (e.g., Saos2, SJSA1, OVSAHO, HCC1937, and HEK293). Unlike **pS1**, the parent phosphopeptide, **pO1**, taking longer time for dephosphorylation than that of **pS1**, requires hours for cellular uptake and largely remains in endosomes. These results indicate that rapid dephosphorylation of the thiophosphate group and the resulting thiol group are critical for instantly targeting Golgi. Based on this insight, we designed **pS2**, a nonfluorescent analogue of **pS1**. Being able to undergo rapid dephosphorylation catalyzed by ALP to form **S2** that exhibit a critical micelle concentration (CMC) of 9.5 μM, **pS2** inhibits HeLa cells with an IC_50_ value about 3 μM, an order of magnitude more potent than that of the parent phosphopeptide. Preliminary mechanistic studies indicate that (i) the thiophosphopeptides enter cells via both caveolin-mediated endocytosis and macropinocytosis, (ii) disulfide bond formation is essential for Golgi targeting, and (iii) the level of ALP of cells contributes to the rate of the accumulation of the resulting thiopeptide assemblies at the Golgi. Providing the first case of targeting Golgi based on fast enzymatic kinetics and oxidative environment of Golgi, this work illustrates a new molecular platform for designing enzyme responsive molecules that target subcellular compartment for functions.

**Scheme 1.**
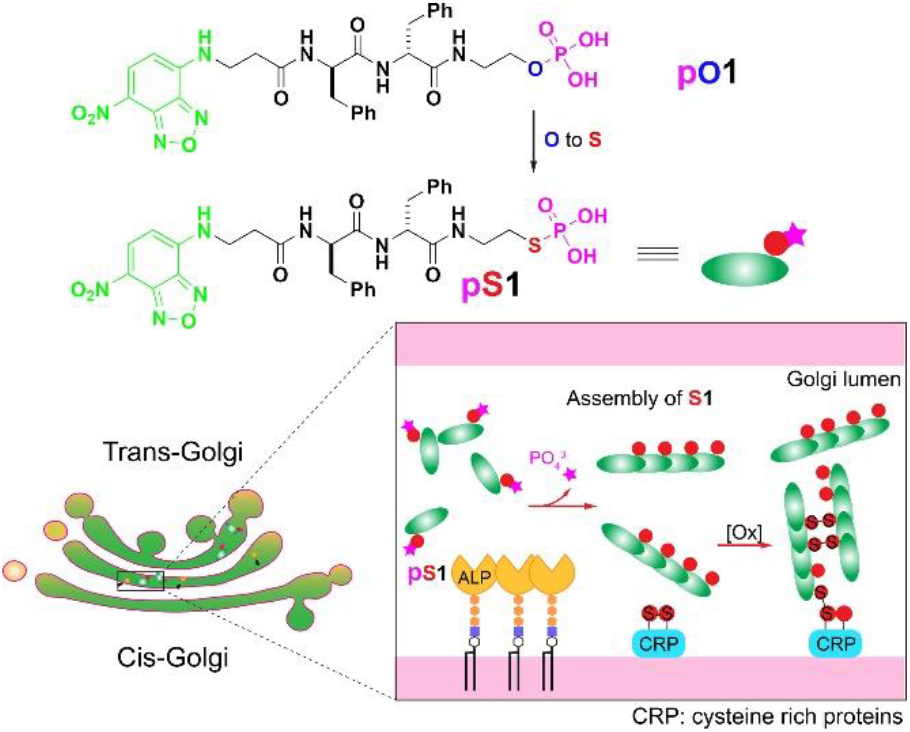
Illustration of thiophosphopeptides instantly targeting the Golgi apparatus.

As shown in Scheme 1, **pS1** consists of three segments: (i) 4-nitro-2,1,3-benzoxadiazole (NBD), a fluorophore that emits bright green fluorescence in hydrophobic environment of supramolecular assemblies;^13^ (ii) D-diphenylalanine (ff), a hydrophobic building block, which enables self-assembly and resists proteolysis; (iii) thiophosphate group, a substrate of ALP^14^ for enzymatic self-assembly.^15-16^ Solid phase peptide synthesis (SPPS) of NBD-ff followed by a conjugation of cysteamine S-phosphate^17^ generates **pS1** (Scheme S2) in a good yield. Such a design ensures the fast dephosphorylation of the thiophosphopeptide by ALP (Figure S1). While **pS1** exhibits critical micelle concentration (CMC) of 6.0 μM, **S1** has the CMC of 2.4 μM (Figure S2). Transmission electron microscopy (TEM) reveals that **pS1**, at 5 μM form micelles, which turn into nanofibers after ALP converts **pS1** to **S1** (Figure S3). The formation of micelles of **pS1** likely facilitates the cellular uptake by caveolin-mediated endocytosis, similar to the cellular uptake of peptide amphiphiles.^18-21^

We first incubated HeLa cells with CellLight® Golgi-RFP^22^ for 24 hours to transfect RFP at the Golgi, then incubated the HeLa cells with **pS1** (10 μM) for 8 minutes (Figure 1). The fluorescence from the assemblies of **S1** overlaps with the red fluorescence from all Golgi-RFP, confirming that **pS1** targets the Golgi of the HeLa cells. The fluorescence of **S1** appears almost instantly after adding **pS1** in the culture of HeLa cells (Figure S4). This rate is, at least an order of magnitude, faster than previously reported probes.^9-11^ The intensity of the fluorescence at the Golgi increases significantly with the time of incubation of **pS1**, about 7 times enhancement from 1 minute to 8 minutes. Except the bright fluorescence at the Golgi and the dim fluorescence at the endoplasmic reticulum (ER), the rest of intracellular and extracellular regions of the HeLa remain dark. This result indicates that enzymatic assembly of **S1** occurs at the Golgi of HeLa cells, agreeing with the observation of ALP at Golgi of HeLa cells.^23^ Live cell imaging over 20 minutes (Video S1) reveals that the fluorescence of the assemblies of **S1** emerges at the Golgi prior to diffusing to ER, likely resulted from Golgi-ER transport.^2^ The differential interference contrast (DIC) image of HeLa cells treated with 10 μM of **pS1** for 20 min (Figure S5) shows well-spread HeLa cells, excluding that **pS1** enters cells due to cell death.

**Figure 1.**
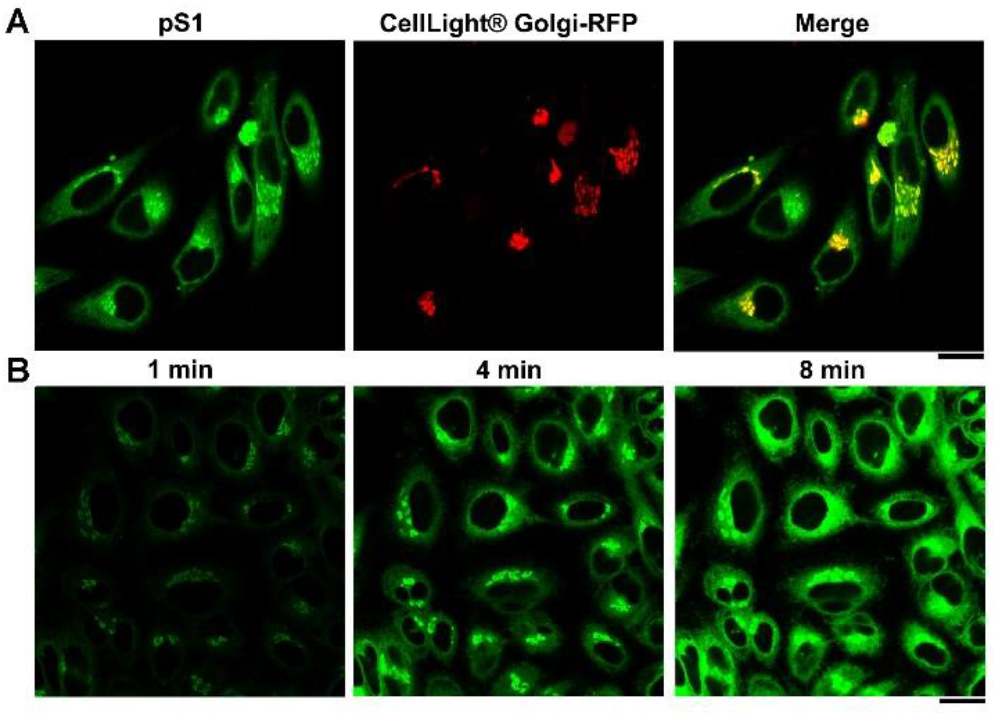
CLSM images of HeLa cells (A) stained with CellLight® Golgi-RFP after treating with **pS1** for 8 minutes and (B) treated with **pS1** for 1, 4, and 8 min. Scale bars = 20 μm and [**pS1**] = 10 μM.

The concentration of **pS1** is another important factor determines the accumulation of **S1** at Golgi. We compared the Golgi fluorescence by fixing the incubation time at 4 minutes and varying the concentration **pS1** at 10, 5, and 2 µM (Figure 2). At 10 μM, bright green fluorescence presents in Golgi and weak fluorescence in ER; at 5 µM, green fluorescence clearly still presents at the Golgi, but little at the ER; at 2 µM, much weaker fluorescence at the Golgi. Further decreasing the concentration of **pS1** to 500 nM still results in Golgi accumulation of **S1** in HeLa cells. Though the brightness of **S1** at Golgi is weaker at the beginning of the addition, distinctive fluorescence appears at the Golgi after 15 minutes (Figure S6). The accumulation rate of **S1** at Golgi, quantified by the increase of the fluorescence intensity, is concentration dependent (Figure S7). Using a Golgi disruptor, brefeldin A (BFA), is able to abolish the accumulation of **S1** (Figure S8). These results indicate that enzymatic formation and self-assembly of **S1** in situ at Golgi enables the instant targeting of Golgi. In addition, the concentration needed of **pS1** for targeting Golgi is much less than the previously reported probes.^9-11^

**Figure 2.**
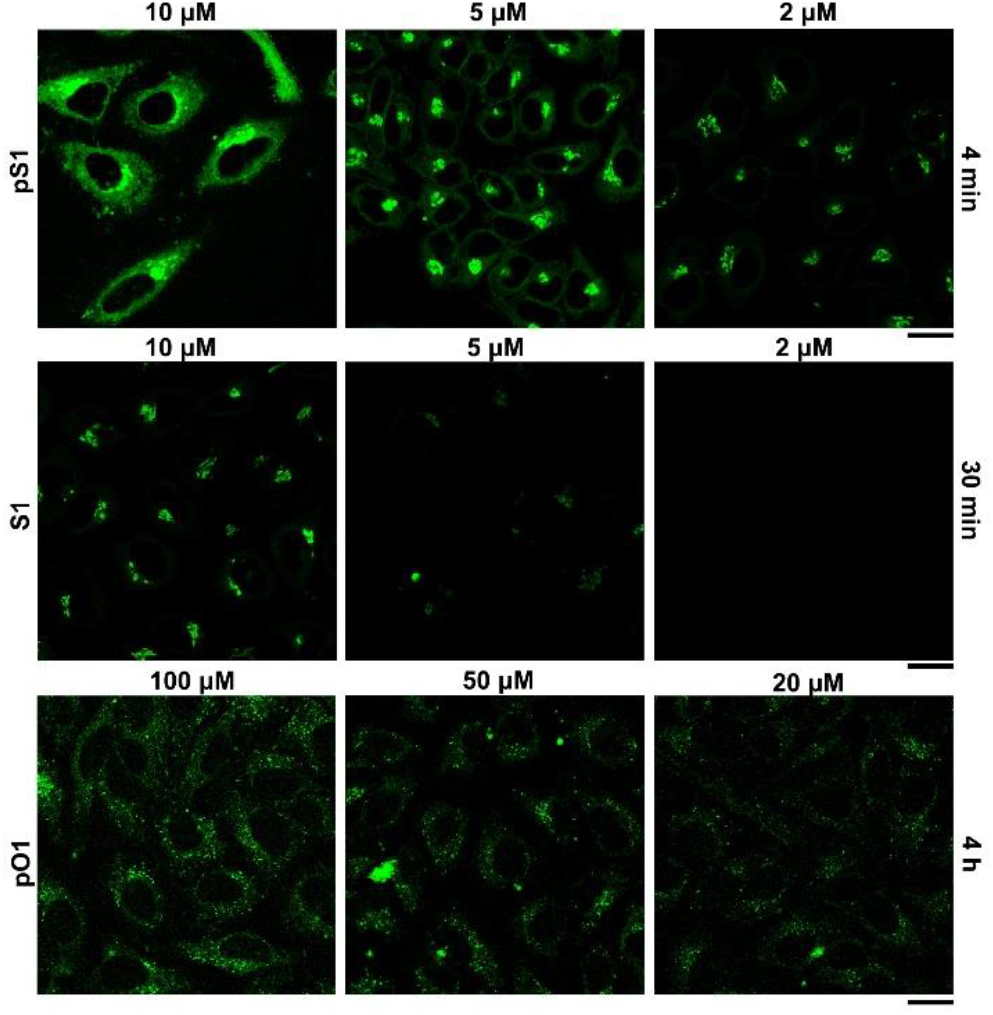
CLSM images of HeLa cells treated with **pS1** (10 μM, 5 μM, 2 μM) for 4 min (top), **S1** (10 μM, 5 μM, 2 μM) for 30 min (middle) and **pO1** (100 μM, 50 μM, 20 μM) for 4 h. Scale bars = 20 μm.

Unlike **pS1, S1**, at the concentration of 10 μM and being incubated with HeLa cells for 8 minutes, hardly results in any fluorescence in the cells (Figure S9A), suggesting slower cell uptake of **S1** than that of **pS1** and indicating the importance of enzymatic dephosphorylation for targeting Golgi. As shown in Figure 2, after incubating **S1** (10 μM) with HeLa cells for 30 minutes, some green fluorescence at the Golgi of the HeLa cells, with the fluorescent intensity similar to that of HeLa cells incubated with **pS1** (2 µM) for 4 minutes. After 30 minutes incubation, when the concentrations of **S1** are at 5 and 2 µM, there is very weak at the Golgi and no fluorescence in cell at all, respectively. These results suggest that **S1** enters the HeLa cells less efficiently than **pS1** does. Moreover, **pO1** (the parent compound of **pS1**) produces almost no fluorescence (Figure S9B) in the HeLa cells after 8 minutes incubation and at the concentration of 10 μM. In fact, after 4 hours of incubation of **pO1** and HeLa cells, there are several scattered fluorescent puncta in cells, with fluorescent intensity being proportional to the concentrations of **pO1** (from 100 to 50 and to 20 µM). These results indicate that the assemblies of **O1**, formed by dephosphorylation, largely are retained in endosomes or lysosomes (Figure 2, Scheme S3). The above results confirm the unique ability of **pS1** for instantly targeting Golgi.

To further understand the mechanism of Golgi-accumulation of **S1** assemblies resulted from the rapid enzymatic dephosphorylation of **pS1**, we examined the rate of fluorescence increase in Golgi for 16 minutes in the HeLa cells treated **pS1** and several inhibitors, using the fluorescence increase in the Golgi of the HeLa cells incubated with **pS1** only as the reference (Figure 3A). Using phospholipase C (PLC),^24^an enzyme that cleaves glycosylphosphatidylinositol (GPI) anchor, to remove ALP from the cell membrane results in slightly faster increase of fluorescence in the Golgi, confirming that ALP at Golgi dephosphorylates **pS1** and indicating that pericellular dephosphorylation of **pS1** by the ALP on plasma membrane slightly slows down the accumulation of **S1** at the Golgi. Methyl-β-cyclodextrin (mβCD) significantly decreases the rate of the fluorescence increase at the Golgi, indicating that **pS1** (at 10 μM) likely enters cells via caveolin-mediated endocytosis. As a potent inhibitor of actin polymerization,^25^ cytochalasin D (CytD) decreases the accumulation of **S1** at Golgi in a concentrationdependent manner (Figure S10). This result agrees with the critical role of actin dynamics in cellular uptake, indicating that **pS1** also enters the cells via marcopinocytosis.^26^ Both the phosphatase inhibitor cocktail set 3 (PIC) and the tissue nonspecific ALP inhibitor (DQB^27^) reduce the rate of the fluorescence increase at the Golgi, with PIC more effectively inhibiting the accumulation than DQB. These results suggest that other phosphatases, in addition to ALP, also contribute to the enzymatic accumulation of **S1** at Golgi and agree with sorting of ALP at the Golgi before secretion.^28-29^

**Figure 3.**
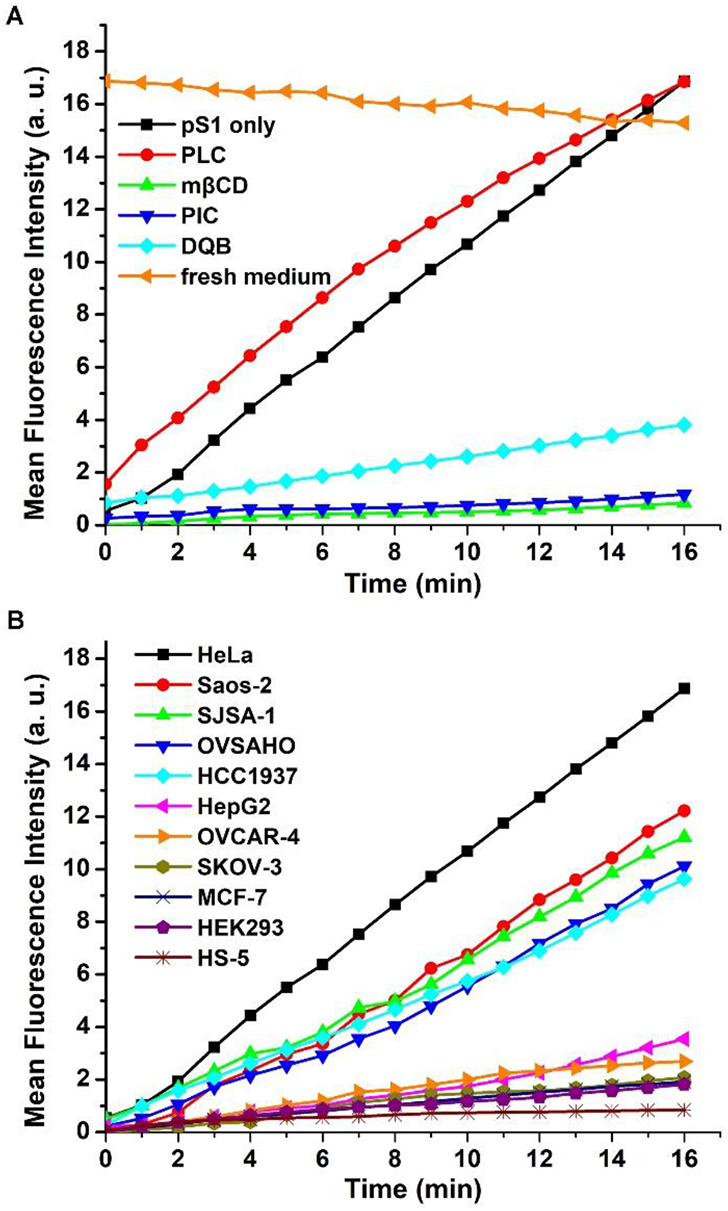
Time-dependent mean fluorescence intensity of Golgi in (A) HeLa cells pretreated with PLC (phospholipase C, 0.2 U, 30 min), mβCD (5 mM, 30 min), PIC (phosphatase inhibitor cocktail set 3, 4000×, 30 min), DQB (20 μM, 30 min), respectively, and then treated with **pS1** (10 μM), and the **pS1** treated HeLa cells in a fresh medium. (B) Time-dependent mean fluorescence intensity of Golgi in different cell lines treated with **pS1** (10 μM).

We also examined the exocytosis of the accumulated **S1** by incubating the HeLa cells pretreated with **pS1** in a fresh culture medium. The intensity of fluorescence at the Golgi of the HeLa cells drops only slightly over time, confirming that the enzymatically formed assemblies of **S1** are largely trapped in the Golgi. Inhibiting protein disulfide isomerases (PDIs) decrease disulfide bonds of cysteine rich proteins (CRPs)^30^ that are transported to Golgi, contributing to a slight decrease of the rate of Golgi accumulation of **S1** (Figure S11). This result implies the peptide assemblies likely form disulfide bonds with CRPs. This observation agrees with that dimers of **S1** (or **S2**) form in the cell lysate treated with **pS1** (or **pS2**) (Figure S12), suggesting that certain extent of covalent linkage between assemblies likely contributes the retention of the assemblies in the Golgi. Moreover, the addition of N-ethylmaleimide almost completely eliminates the accumulation of **S1** at Golgi (Figure S13), further supporting that **S1** forms disulfide bond with CRPs at Golgi.

To examine the applicability of the process illustrated in Scheme 1 for targeting Golgi of other cells, we incubated **pS1** with several other cancer cell lines (Saos-2, SJSA-1, OVSAHO, HCC1937, HepG2, OVCAR-4, SKOV-3, MCF-7) and immortalized normal cell lines (HEK293 and HS-5) and examined the rates of fluorescent increase at the Golgi of the cells (Figure 3B). The fluorescence intensities increase significantly at the Golgi of Saos-2, SJSA-1, OVSAHO and HCC1937 cells, slightly in those of HepG2 and OVCAR4 cells, and much slowly in those of SKOV3, MCF7, HEK293 and HS-5 cells. These results largely agree with expression levels of ALP in these cell (Figure S14).^31^ One exception is HepG2, which expresses higher level of ALP than OVSAHO, but exhibits slower fluorescence increase at Golgi. High level of glutathione in hepatocytes^32^ likely antagonizes the accumulation of **S1** in the Golgi. This observation supports that oxidative Golgi environments favors disulfide formation, thus the retention of the assemblies of **S1** at the Golgi.

We synthesized a nonfluorescent analogue (**pS2**) of **pS1** by using naphthyl group to replace NBD (Figure S15). Being similar as **pS1, pS2** undergoes rapid dephosphorylation by ALP to form **S2** (Figure S16). Compared with its oxophosphate analogue **pO2**,^33^ **pS2** shows much faster dephosphorylation. For example, being incubated with ALP (0.1 U/mL) for about 16 minutes, **pS2** nearly fully converted to **S2**, while the maximum conversion ratio of **pO2** to **O2** remains at about 50% at the same duration (Figure S16). The CMC of **pS2** is 9.5 μM, and the CMC of its dephosphorylated product, **S2**, is 4.3 μM (Figure S17), indicating both compounds have an excellent self-assembling ability. We further tested the cytotoxicity of **pS2** against HeLa, HEK293, and HS-5 cells and found the IC50 values are 2.8 μM, >100 μM and >100 μM, respectively (Figure 4A). The inhibitory activity of **pS2** against HeLa cells is an order of magnitude higher than that of **pO2** (Figure S18). The difference between these IC50 values agrees with the difference of the rate of enzymatic assemblies in Golgi of the cells, indicating that selectively targeting the Golgi is resulted from fast enzyme kinetics. This result also confirms that **pS2** is more selective than **S2** against cancer cells (Figure S19). In addition, several commonly used inhibitors (Z-VAD-FMK, NAc, Nec-1, DFO, Fer-1, and disulfiram)^34^ of cell death are unable to rescuing HeLa cells from **pS2** (Figures S20 and S21), suggesting a unique cell death resulted from the molecular processes defined thiophosphopeptides at the Golgi apparatus. Treating the HeLa cells incubated with **pS2** by the tetracysteine probe, FlAsH-EDT2,35 results in the fluorescent at the Golgi of the HeLa cells (shown by the arrow in Figures 4B and S22), indicating that **S2** self-assembles at the Golgi to arrange multiple C-terminal thiols into arrangement similar to that of tetracysteine.

**Figure 4.**
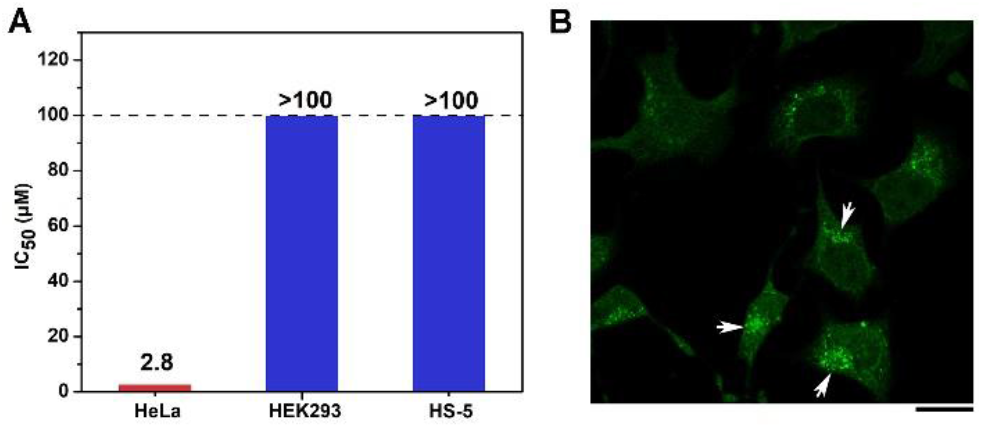
(A) IC50 of **pS2** against HeLa cells, HEK293 cells and HS-5 cells. (B) CLSM images of HeLa cells treated with **pS2** (10 μM, 4 h) and then stained by a tetracysteine probe, FlAsH-EDT2. Scale bar = 20 μm.

In summary, this work illustrates that rapid dephosphorylation of thiophosphopeptides enables instantly targeting of Golgi apparatus and selectively inhibiting the cancer cells. Our observations agree with several known facts: (i) CRPs are enriched in Golgi,^36-38^ (ii) a significant level of oxidation occurs in the Golgi membrane,^39^ (iii) ALP, as an “almost perfect” enzyme^40^ being anchored on the cell membrane by glycosylphosphatidylinositol (GPI) and overexpressed on certain cancer cell,^23, 41-42^ is known to be sorted as oligomers at the Golgi before secretion.^28-29^ Since thiophosphopeptides^43-45^ are much less developed than phosphopeptides, replace NBD with other functional motifs (*e*.*g*. 10-hydroxycamptothecine)^46^ may lead to new discoveries. These thiophosphopeptides also may serve as the substrates for thiol-click chemistry^47^ or for integration thiol groups in other supramolecular assemblies.^48-52^

## ASSOCIATED CONTENT

### Notes

The authors declare no competing financial interests.

## ACKNOWLEDGMENT

This work is partially supported by NIH (CA142746, CA252364) and NSF (DMR-2011846).

---

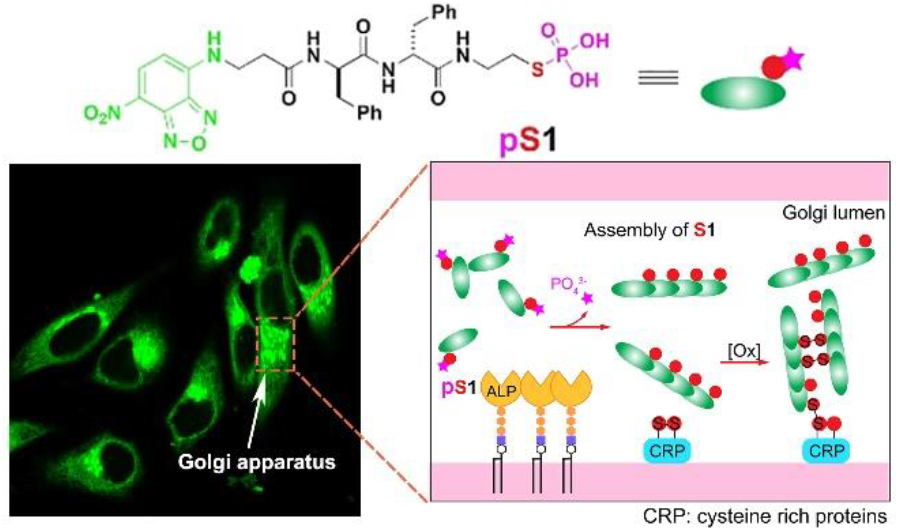

