## Supplementary Information for "Thiophosphopeptides Instantly Targeting Golgi Apparatus and Selectively Killing Cancer Cells"

**Content**

S1. Experiment materials and instruments……………………………………..…S4

S2. Synthesis and characterization of the compounds………………………..……S5

**Scheme S1.** Synthetic procedure of NBD-β-Alanine.

**Scheme S2.** Synthetic procedure of **pS1** and **pS2**.

S3. Cell culture…………………………………………………….………………S7

S4. Critical micelle concentration (CMC) determination…………………….....S7

S5. TEM experiments………………………………………………….………....S8

S6. Determination of dephosphorylation rates *in vitro*…….…………………….S8

S7. Staining cells with FlAsH-$EDT_2$…………………………………...…………S8

S8. Supplemental figures………………………….…………………….……….. S8

**Scheme S3.** Illustration of slow enzymatic reactions of **pO1** to go into cells and chemical structures of **pO1** and **O1**.

**Figure S1.** Time-dependent dephosphorylation of **pS1** (120 μM) and **pO1**

(120 μM) treated with ALP.

**Figure S2.** CMC determination of **pS1** and **S1** by dynamic light scattering (DLS).

**Figure S3.** TEM images of **pS1** with or without the addition of ALP for 24 h.

**Figure S4.** CLSM image of HeLa cells at 0 min immediately after the treatment of **pS1** (10 μM).

**Figure S5.** DIC image of HeLa cells treated with **pS1** (10 μM) for 20 min.

**Figure S6.** CLSM and DIC image of HeLa cells treated with **pS1** (500 nM) for 15 min.

**Figure S7.** Time-dependent mean fluorescence intensity of HeLa cells which are treated different concentrations of **pS1**.

**Figure S8.** CLSM and DIC image of HeLa cells pretreated with BFA and then treated with **pS1** (10 μM) in 16 min.

**Figure S9.** CLSM images of HeLa cells treated with **S1** (10 μM) and treated with **pO1** (10 μM).

**Figure S10.** Time dependent mean fluorescence intensity of Golgi in HeLa cells treated with only **pS1**, and pretreated with different concentrations of CytD (0.5 μg/mL, 1.0 μg/mL) for 30 minutes and then treated with **pS1** (10 μM).

**Figure S11.** Time dependent mean fluorescence intensity of Golgi in HeLa cells treated with only **pS1**, and pretreated with PDI inhibitor (PACMA31, 10 μM) for 30 minutes and then treated with **pS1** (10 μM).

**Figure S12.** LC/MS spectrum of (A) **pS1** and (B) **pS2** treated with HeLa cell lysate for 24 h at 37 ℃.

**Figure S13.** CLSM images of HeLa cells treated with or without NEM (5 mM) combining with **pS1** (10 μM) for 16 min.

**Figure S14.** Accumulative ALP expression levels and the apparent rate of the increase of the fluorescence at the Golgi of different cell lines.

**Figure S15.** Structures of **pS2** and **S2**.

**Figure S16.** Time-dependent dephosphorylation of **pS2** (120 μM) and **pO2** (120 μM) treated with ALP.

**Figure S17.** CMC determination of **pS2** and **S2** by dynamic light scattering (DLS).

**Figure S18.** Cytotoxicity of **pS2** and **pO2** against HeLa cells for 24 h and the values of $IC_{50}$ for both phosphopeptides.

**Figure S19.** Cytotoxicity of **pS2** and **S2** against HeLa cells, HEK293 cells and HS-5 cells.

**Figure S20.** Cell viability of HeLa cells treated with only **pS2** and the mixture of **pS2** with apoptosis and necroptosis inhibitors (Z-VAD-FMK; NAc; Nec-1) for 24 h.

**Figure S21.** Cell viability of HeLa cells treated with only **pS2** (10 μM), the mixture of **pS2** (10 μM) with ferroptosis inhibitors (DFO; Fer-1) and pyroptosis inhibitor (Disulfiram) for 24 h.

**Figure S22.** CLSM of HeLa cells with or without the pretreatment of **pS2** (10 μM, 30 min) and then with or without staining by FlAsH-EDT$_2$ (1 μM, 1 h).

**Figure S23.** LC/MS data for **pS1**.

**Figure S24.** LC/MS data for **pS2**.

**Figure S25.** LC/MS data for **pO1**.

**Figure S26.** LC/MS data for **pO2**.

**Video S1.** CLSM of HeLa cells treated with **pS1** (10 μM) for 16 min.

### S1. Experiment materials and instruments

2-Cl-trityl chloride resin (1.0 mmol/g), Fmoc protected amino acid, and HBTU were obtained from GL Biochem (Shanghai, China). N, N-diisopropylethylamine (DIEA) and solvents were obtained from Fisher Scientific. Alkaline phosphatase was purchased from Biomatik (Cat. No. A1130, alkaline phosphatase [ALP], >1300U/mg, in 50% glycerol.). Cysteamine S-phosphate, cysteamine hydrochloride and O-phosphorylethanolamine were all purchased from Sigma-Aldrich. All the chemical reagents and solvents were used as received from commercial sources without further purification. Minimum Essential Media (MEM), Dulbecco's Modified Eagle Medium (DMEM), McCoy's 5A Medium, and RPMI-1640 Medium were purchased from ATCC. Fetal bovine serum (FBS) and Penicillin-Streptomycin from Gibco by Life Technologies. All precursors and compounds were purified by a reverse phase HPLC (Agilent 1100 Series) equipped with an XTerra C18 RP column, and HPLC grade acetonitrile (0.1% TFA) and HPLC grade water (0.1% TFA) were used as the eluents. The LC-MS spectra were obtained with a Waters Acquity Ultra Performance LC with Waters MICROMASS detector. Transmission electron microscope (TEM) images were obtained on Morgagni 268 transmission electron microscope. 1H-NMR spectra of compounds were obtained using Varian Unity Inova 400MHz. Fluorescence images were taken by ZEISS LSM 880 confocal laser scanning microscope.

### S2. Synthesis and characterization of the compounds

### Synthesis of NBD-β-Alanine

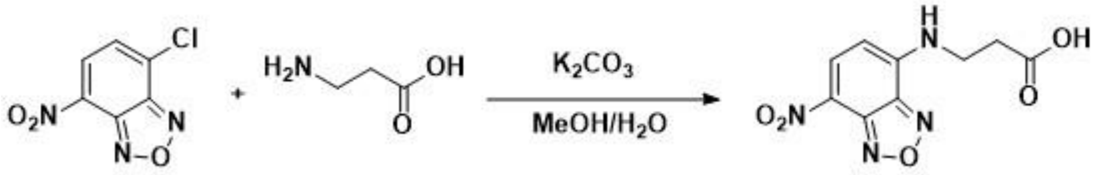


**Scheme S1.** Synthetic procedure of NBD-β-Alanine.

To a 10 mL water solution of β-Alanine (5.5 mmol, 490 mg) and potassium carbonate (16.5 mmol, 2.07g), NBD-Cl (5 mmol, 1g) in 60 mL of MeOH was added to the above solution dropwise with stirring under nitrogen gas protection. After stirring at room temperature for 6 h, methanol was removed by a rotary evaporator and acidified the residual solution to pH 3 by HCl (1 N). The acidic aqueous solution was then extracted by diethyl ether. The combined organic solution was dried over anhydrous sodium sulfate, and then concentrated by a rotary evaporator. The resulting dark-yellow powder (NBD-β-Alanine) was directly used for solid phase peptide synthesis.

### Peptide synthesis

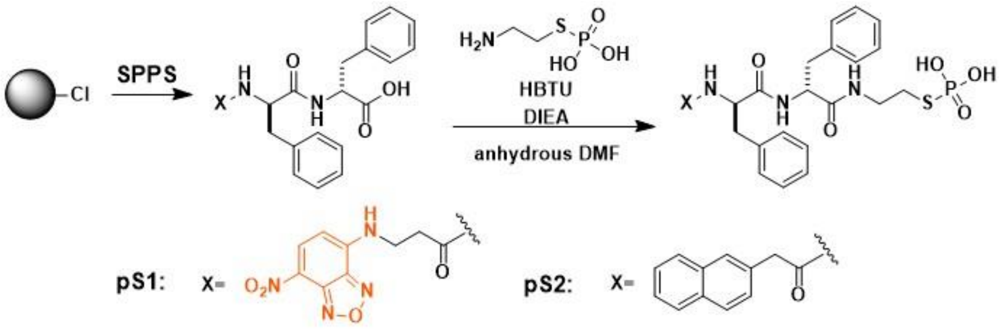


**Scheme S2.** Synthetic procedure of **pS1** and **pS2**.

#### Solid Phase Peptide Synthesis

We used standard Fmoc chemistry solid phase peptide synthesis using 2-chlorotrityl chloride resin and the corresponding Fmoc-protected amino acids with side chains properly protected.

Briefly, the 2-Cl resin (1 g) was swelling in dry DCM for 30 min and then the first amino acid was loaded onto the resin. After loading the first amino acid to the resin, the capping regent (DCM: MeOH: DIEA = 17: 2: 1) was used to cap all the active sites of the resin for additional 15 min. Fmoc group was then removed with 20% piperidine in DMF, the next Fmoc-protected amino acid was coupled to the free amino group using HBTU as the coupling reagent. The peptide chain was cleaved from the resin by 95% TFA (95% TFA, 2.5% TIPS, 2.5% $H_2O$) for 1 h. After the solvent being removed by a rotary evaporator, 30 mL of dry diethyl ether was added to the residual solution and followed by centrifuging at 10000 rpm for 8 min. The resulting solid products were dried by a lyophilizer. And we got a compound in solid orange powder (NBDff) and a compound in solid white powder (Napff).

**Synthesis of pS1 and pS2**

Briefly, 0.1 mmol of synthesized peptide (NBDff or Napff) was dissolved in 2 mL dry DMF and 0.12 mmol of HBTU was added directly into the solution. After stirring the mixture for 30 minutes, 0.12 mmol of Cysteamine S-Phosphate sodium salt was added into the mixture and DIEA was added dropwise to adjust pH around 8, then keep stirring for 8 h. After that, the solvent is air dried and the remained oily product was dissolved in methanol and purified by RP-HPLC to obtain the titled compounds.

**Synthesis of S1**

Briefly, 0.1 mmol of synthesized peptide (NBDff) was dissolved in 2 mL dry DMF and 0.12 mmol of HBTU was added directly into the solution. After stirring the mixture for 30 minutes, 0.12 mmol of cysteamine hydrochloride was added into the mixture and DIEA was added dropwise to adjust pH around 8, then keep stirring for 8 h. After that, the solvent is air dried and the remained oily product was dissolved in methanol and purified by RP-HPLC to obtain the titled compound.

**Synthesis of pO1 and pO2**

Briefly, 0.1 mmol of synthesized peptide (NBDff or Napff) was dissolved in 2 mL dry DMF and 0.12 mmol of HBTU was added directly into the solution. After stirring the mixture for 30 minutes, 0.12 mmol of O-Phosphorylethanolamine was added into the mixture and DIEA was added dropwise to adjust pH around 8, then keep stirring for 8 h. After that, the solvent is air dried and the remained oily product was dissolved in methanol and purified by RP-HPLC to obtain the titled compounds.

### S3. Cell culture

All cell lines used (HeLa, Saos-2, SJSA-1, OVSAHO, HCC1937, HepG2, OVCAR-4, SKOV-3, MCF-7, HEK293, HS-5) were purchased from the American Type Culture Collection (ATCC, Manassas, VA, USA). HeLa, HepG2, MCF-7 and HEK293 cells were cultured in MEM medium with 10% FBS and 1% P/S (100 U $mL^{-1}$ penicillin and 100 μg $mL^{-1}$ streptomycin, Invitrogen Life Technologies). Saos-2, SKOV-3 cells were cultured in McCoy's 5A medium with 15% FBS and 1% P/S. HS-5 cells were cultured in DMEM medium with 10% FBS and 1% P/S. SJSA-1, OVSAHO, HCC1937, OVCAR-4 cells were cultured in RPMI-1640 medium with 10% FBS and 1% P/S. All the cells were cultured at 37 °C in a humidified atmosphere of 5% $CO_2$. To determine the cytotoxicity of the compounds, cells were seeded in 96-well cell plate at $1.0\times10^4$ cells/well for 24 h followed by culture medium removal, and then fresh culture medium containing compounds were added. For fluorescence imaging, cells were seeded in confocal dish at $1.5\times10^5$ cells/dish for 24 h followed by the addition of fresh medium containing compounds, and then the fluorescence of cells were taken by ZEISS LSM 880 confocal laser scanning microscope with the 63× oil lens.

### S4. Critical micelle concentration (CMC) determination

A series of solutions of compounds (**pS1**, **pS2**), from the concentration of 0.0625 μM to 400 μM, were prepared in deionized water and adjusted the pH to 7.4. The count rates of the solutions were measured and recorded by an ALV/DLS/SLS-5000 Light Scattering System, which were converted to the intensity of scattered light and then plotted against concentrations to determine the CMC values. Then, ALP (0.1 U/mL) was added to the series of solutions and after 24 h, the same procedures were proceeded to determine CMC values of dephosphorylated **pS1** and **pS2** (**S1** and **S2**).

### S5. TEM experiments

The 400 mesh copper grids coated with carbon film was glowing discharged and sample solutions (5 μL of each) was placed onto the grids. After 30 seconds, the sample solution was removed, and the grids were stained by uranyl acetate (2% v/v) and allowed to dry in air. TEM images were obtained with Morgagni 268 transmission electron microscope at the HV of 80 kV with filament of 2.

### S6. Determination of dephosphorylation rates *in vitro*

To determine the dephosphorylation rates of thiophosphopeptides *in vitro*, we prepared **pS1**, **pO1**, **pS2**, **pO2** solution (PBS, pH 7.4) at concentration of 120 μM and ALP was added into the solutions to make the final concentration of ALP to 0.1 U/mL at 37 °C. At the designated time, we added same volume of methanol to eliminate the activity of ALP and used LC/MS to analysis the results.

### S7. Staining cells with FlAsH-EDT$_2$

Briefly, after incubating HeLa cell lines ($1.5\times10^5$) in 3.5 cm confocal dish for 24 h at 37 °C in a humidified atmosphere of 5% $CO_2$, we discarded the medium and added fresh medium containing **pS2** (10 μM) and incubated them together for 4 h. After 4 h, medium was removed followed by three-time wash by cell imaging buffer to eliminate FBS. Then, we added FlAsH-EDT$_2$ solution (cell imaging buffer, 1 μM) into the confocal dish and incubated the HeLa cells at 37 °C for 1 h. Meanwhile, we prepared 1,2-Ethanedithiol (EDT) solution (cell imaging buffer, 250 μM) and after the one-hour incubation of HeLa cells with FlAsH-EDT$_2$, FlAsH-EDT$_2$ was discarded and EDT solution was added to the confocal dish to eliminate the unspecific binding of FlAsH-EDT$_2$ for 10 min. Ten minutes later, EDT solution was removed and cells were washed with cell imaging buffer three times and the fluorescence images were taken by ZEISS LSM 880 confocal laser scanning microscope.

### S8. Supplemental figures

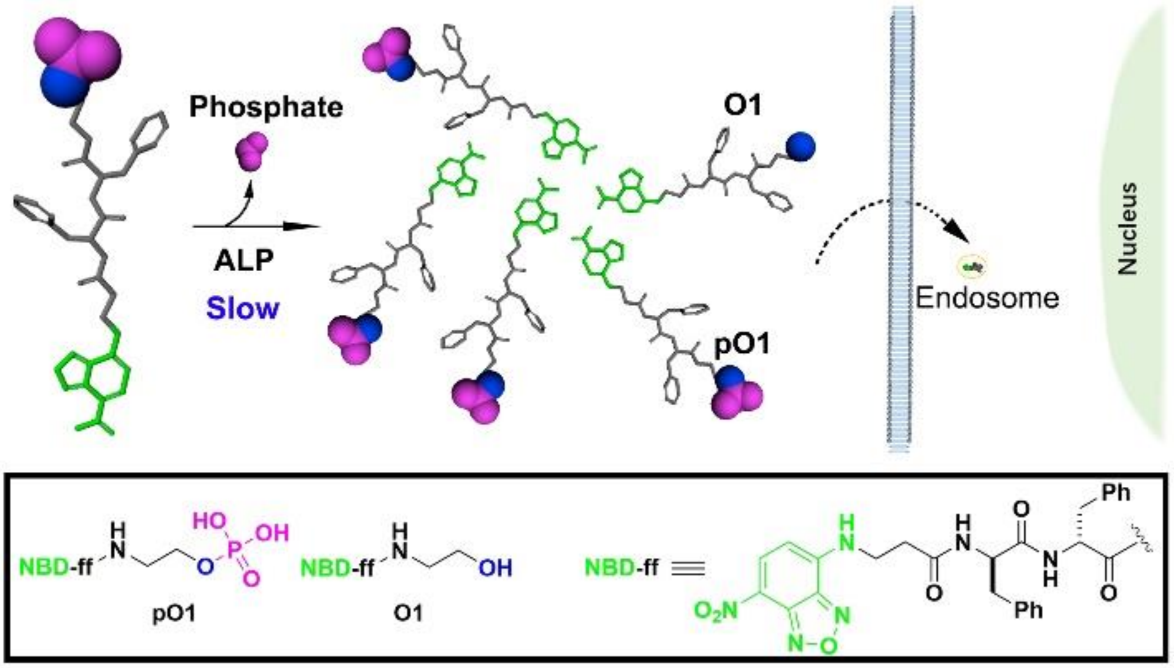


**Scheme S3.** Illustration of slow enzymatic reactions of **pO1** to go into cells and chemical structures of **pO1** and **O1**.

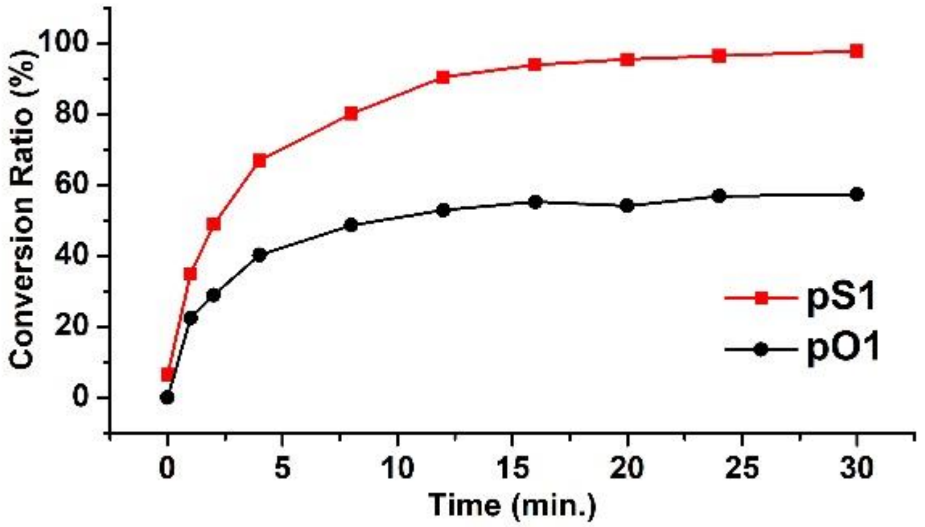


**Figure S1.** Time-dependent dephosphorylation of **pS1** (120 μM) and **pO1** (120 μM) treated with ALP (0.1 U/mL), respectively.

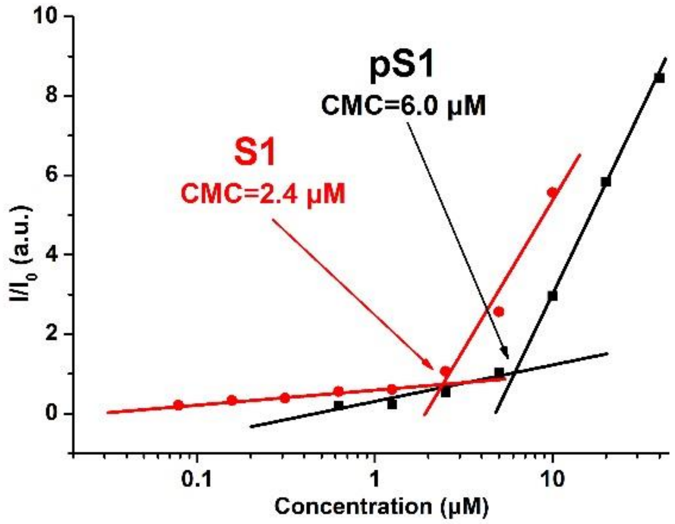


**Figure S2.** CMC determination of **pS1** and **S1** by dynamic light scattering (DLS).

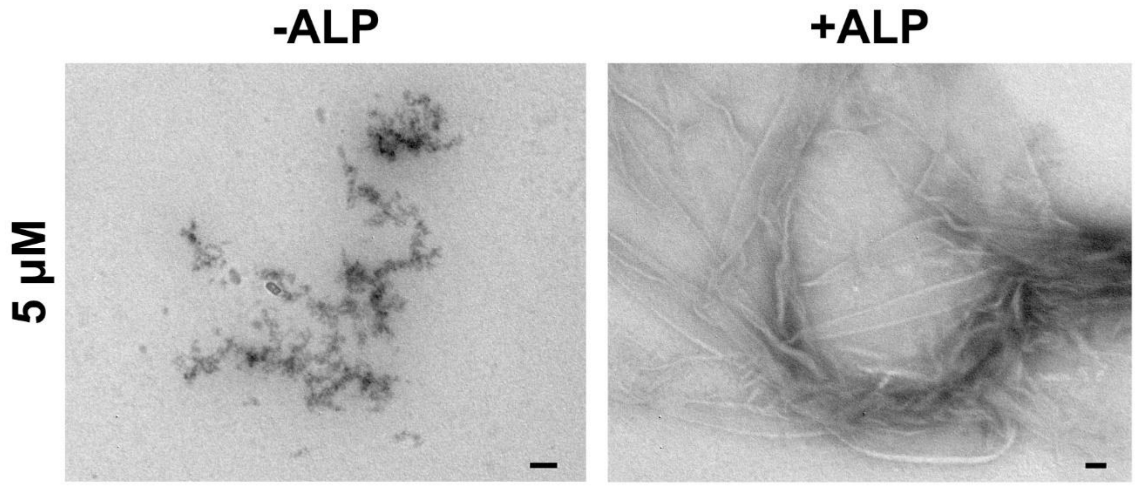


**Figure S3.** TEM images of **pS1** (5 μM) with or without the addition of ALP (0.1 U/mL) for 24 h, Scale bars = 100 nm.

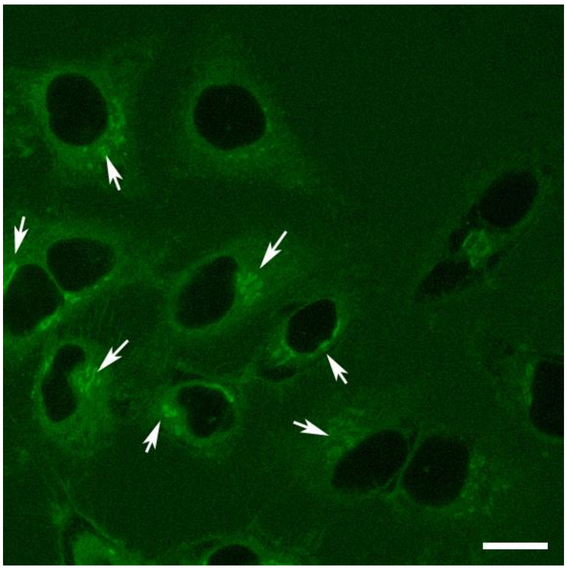

**Figure S4.** CLSM image of HeLa cells at 0 min immediately after the treatment of **pS1** (10 μM). The Golgi of HeLa is marked by arrows. Scale bar = 20 μm.

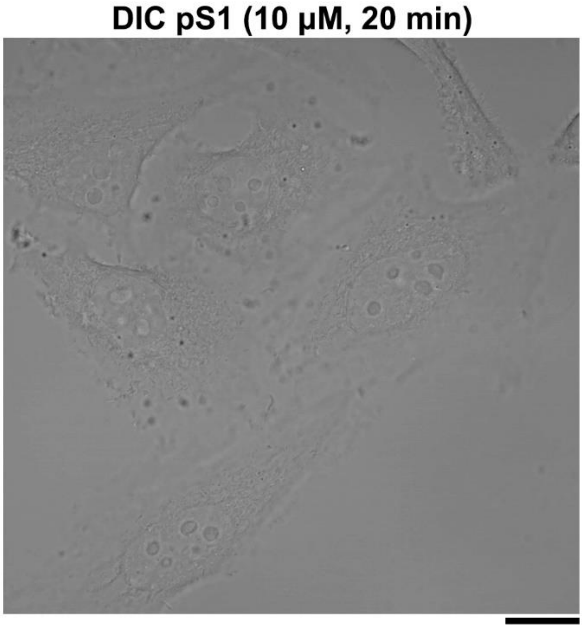


**Figure S5.** DIC image of HeLa cells treated with **pS1** (10 μM) for 20 min. Scale bar = 20 μm.

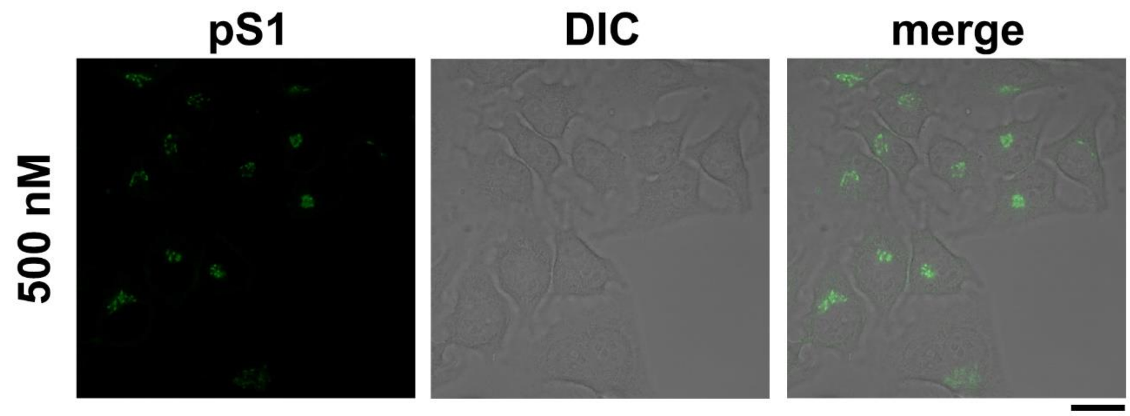


**Figure S6.** CLSM and DIC image of HeLa cells treated with **pS1** (500 nM) for 15 min. Scale bar = 20 μm.

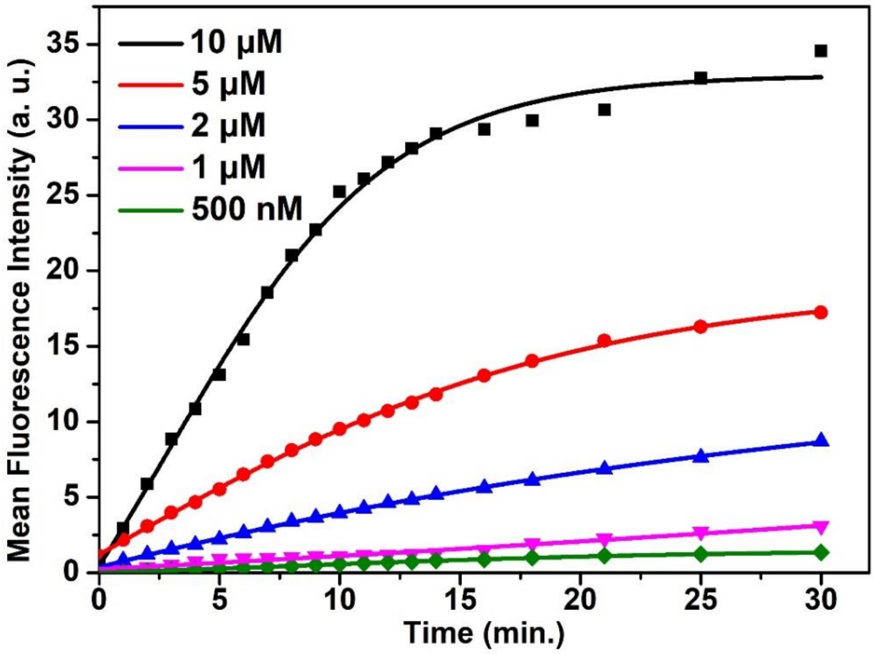


**Figure S7.** Time-dependent mean fluorescence intensity of HeLa cells treated with different concentrations of **pS1**.

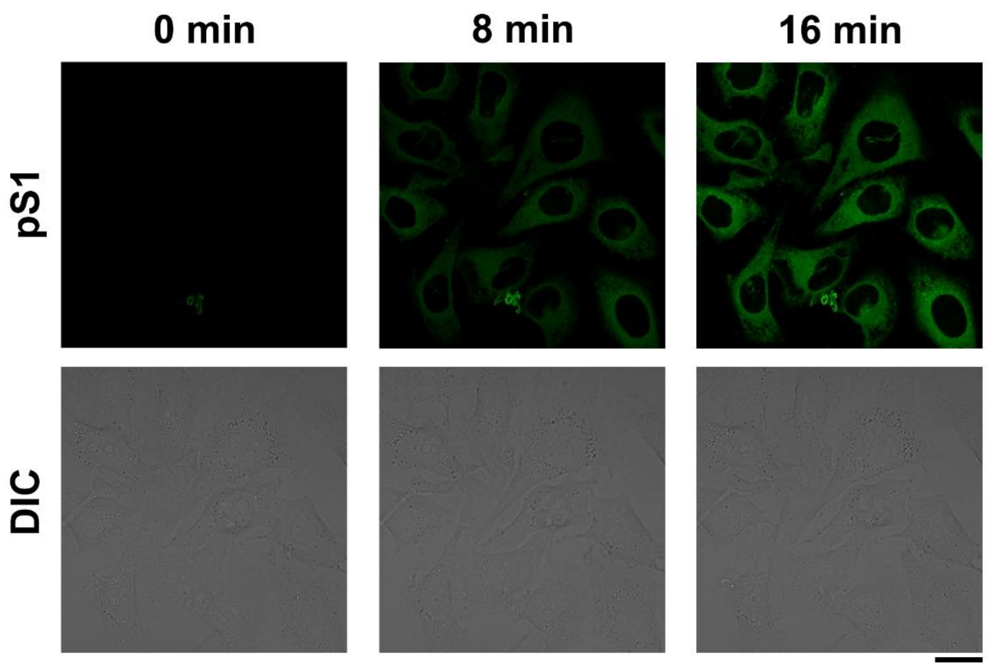


**Figure S8.** CLSM and DIC image of HeLa cells pretreated with BFA (20 μM) for 30 min and then treated with **pS1** (10 μM) in 16 min.

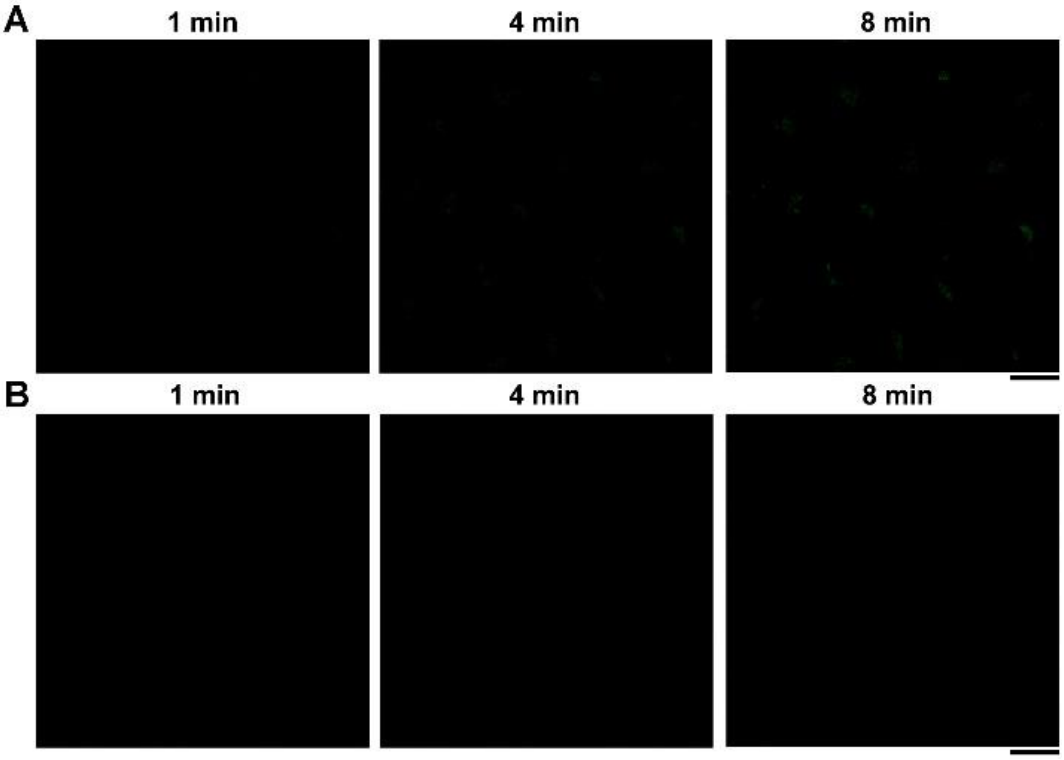


**Figure S9.** CLSM images of HeLa cells (A) treated with **S1** (10 μM) for 1 min., 4min. and 8 min. (B) treated with **pO1** (10 μM) for 1 min., 4min. and 8 min. Scale bars = 20 μm.

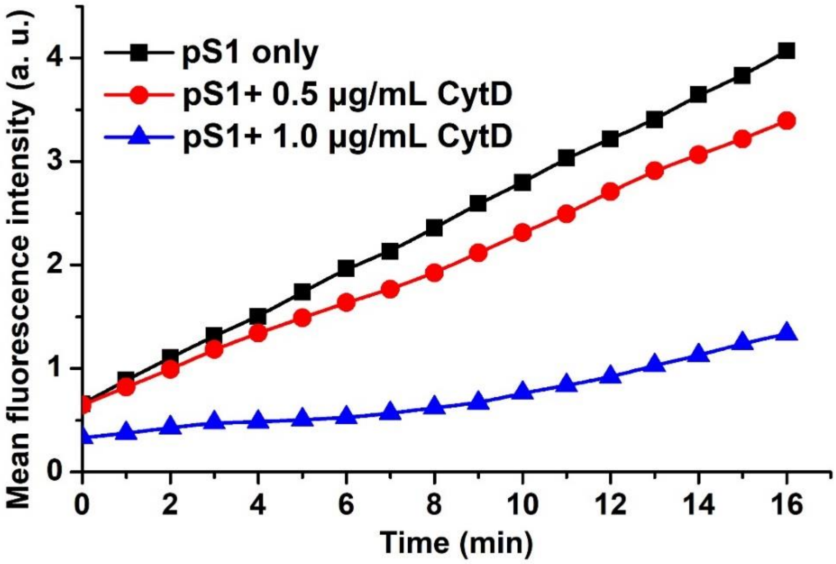


**Figure S10.** Time dependent mean fluorescence intensity of Golgi in HeLa cells treated with only **pS1** (10 μM), and pretreated with different concentrations of CytD (0.5 μg/mL, 1.0 μg/mL) for 30 minutes and then treated with **pS1** (10 μM).

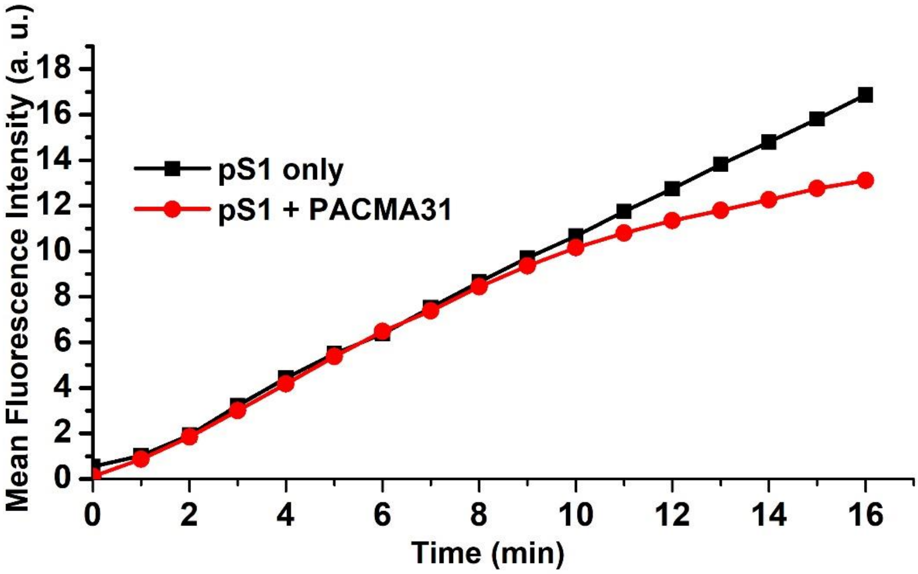


**Figure S11.** Time dependent mean fluorescence intensity of Golgi in HeLa cells treated with only **pS1** (10 μM), and pretreated with PDI inhibitor (PACMA31, 10 μM) for 30 minutes and then treated with **pS1** (10 μM).

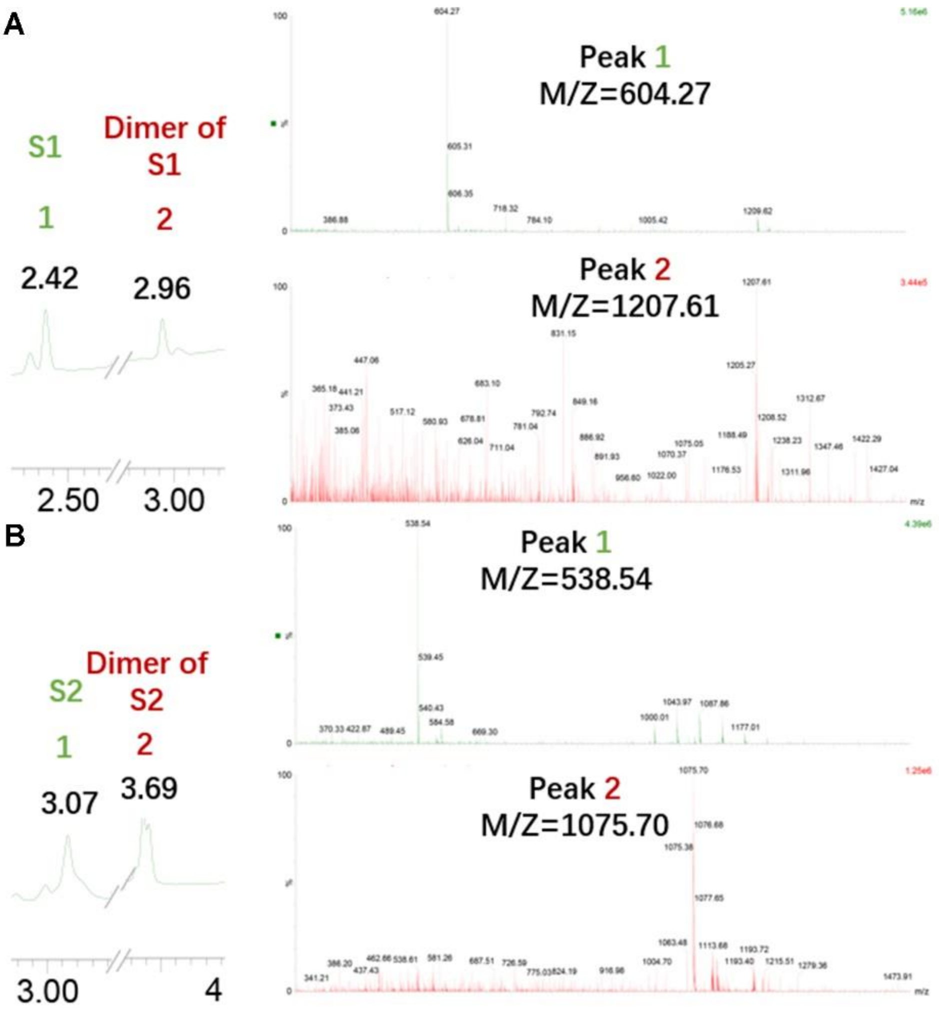


**Figure S12.** LC/MS spectrum of (A) **pS1** and (B) **pS2** treated with HeLa cell lysate for 24 h at 37 ℃. The calculated molecular weight (Mw) of **S1** is 605.21. The observed M/Z =604.27; The calculated molecular weight (Mw) of dimer of **S1** is 1028.40. The observed M/Z =1027.61; The calculated molecular weight (Mw) of **S2** is 539.22. The observed M/Z =538.54; The calculated molecular weight (Mw) of dimer of **S2** is 1076.43. The observed M/Z =1075.70.

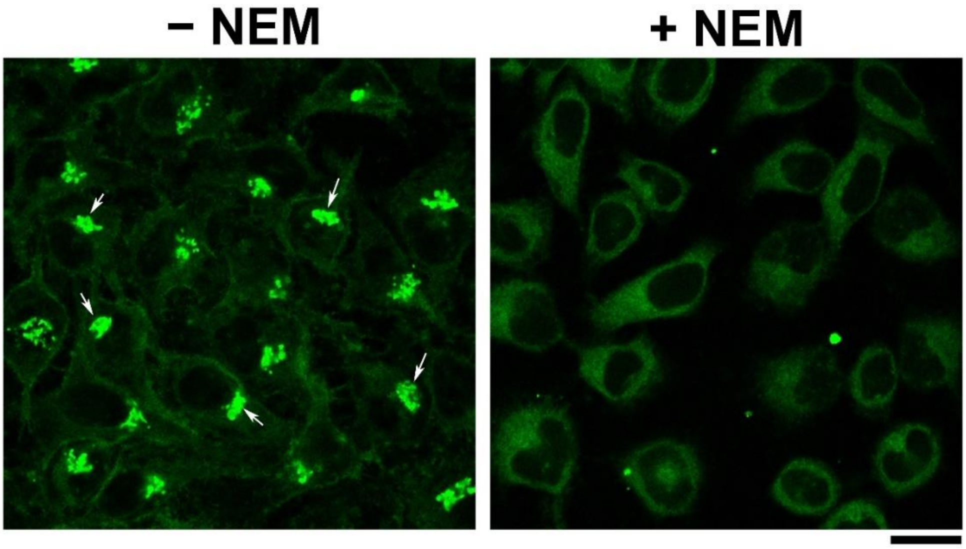


**Figure S13.** CLSM images of HeLa cells treated with or without NEM (5 mM) combining with **pS1** (10 μM) for 16 min. The Golgi of HeLa is marked by arrows. Scale bar = 20 μm.

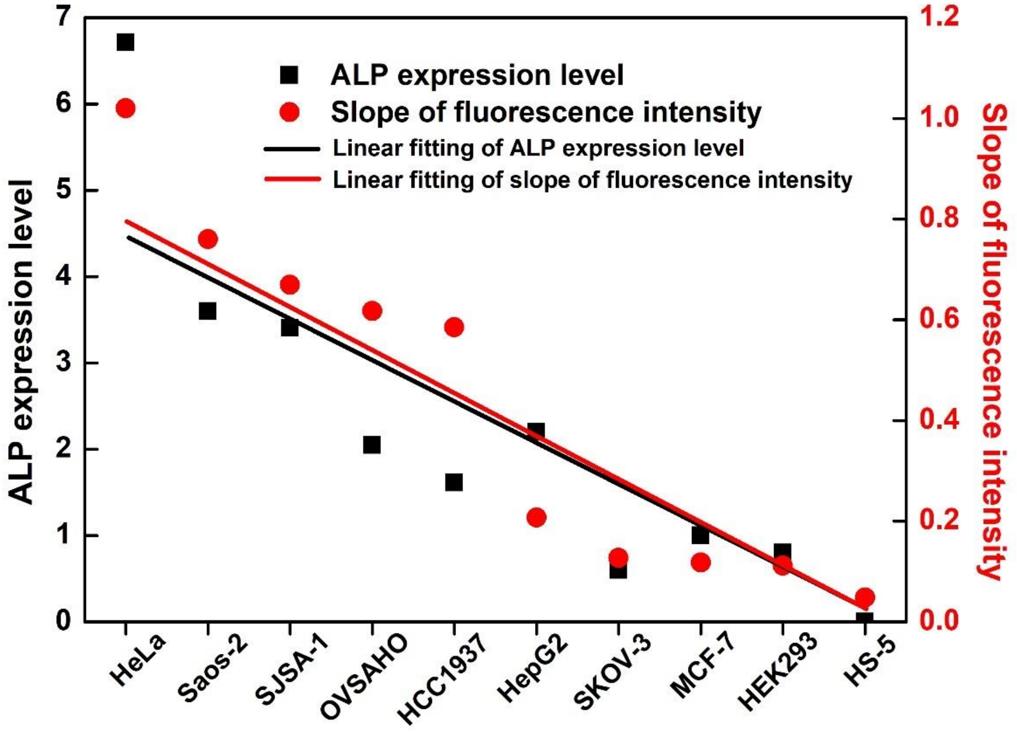


**Figure S14.** Accumulative ALP expression levels (obtained from Harmonizome Database[1]) and the apparent rate of the increase of the fluorescence at the Golgi of different cell lines.

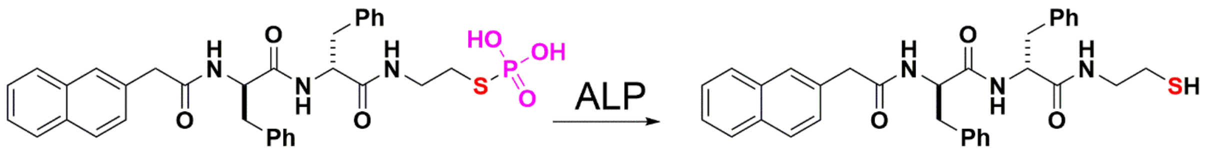


**Figure S15.** Structures of **pS2** and **S2**.

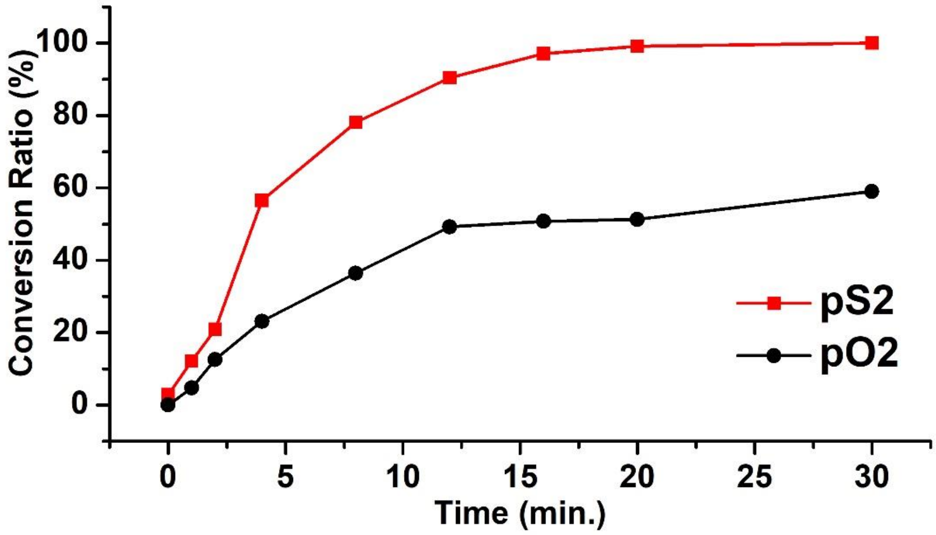


**Figure S16.** Time-dependent dephosphorylation of **pS2** (120 μM) and **pO2** (120 μM) treated with ALP (0.1 U/mL), respectively.

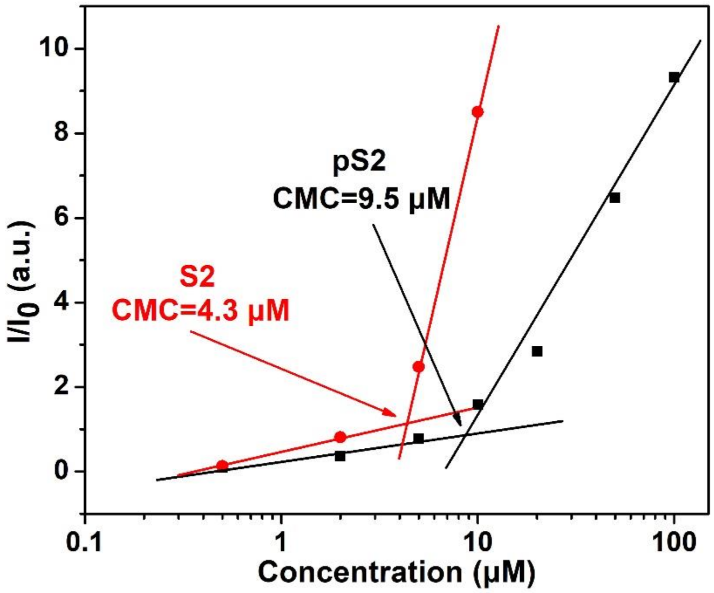


**Figure S17.** CMC determination of **pS2** and **S2** by dynamic light scattering (DLS).

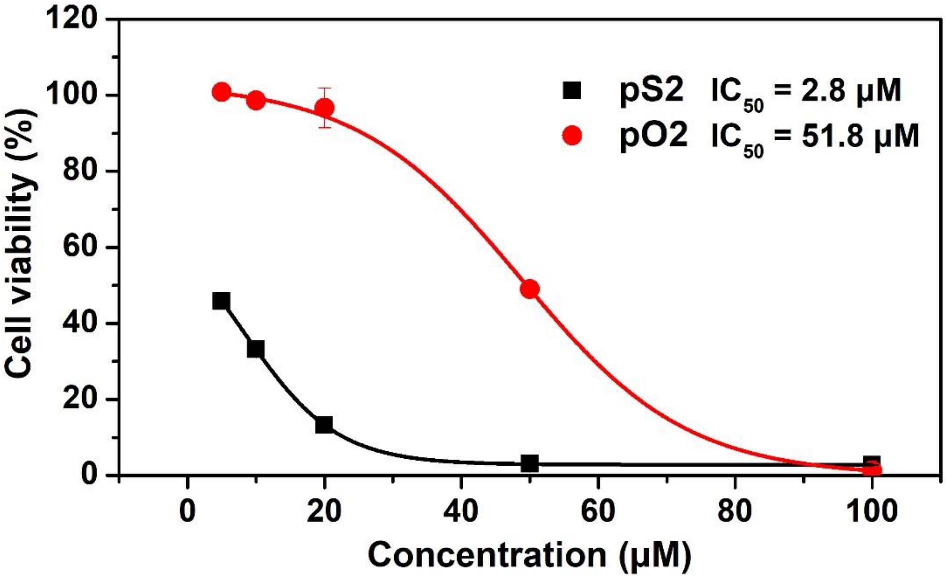


**Figure S18.** Cytotoxicity of **pS2** and **pO2** against HeLa cells for 24 h and the values of $IC_{50}$ for both phosphopeptides.

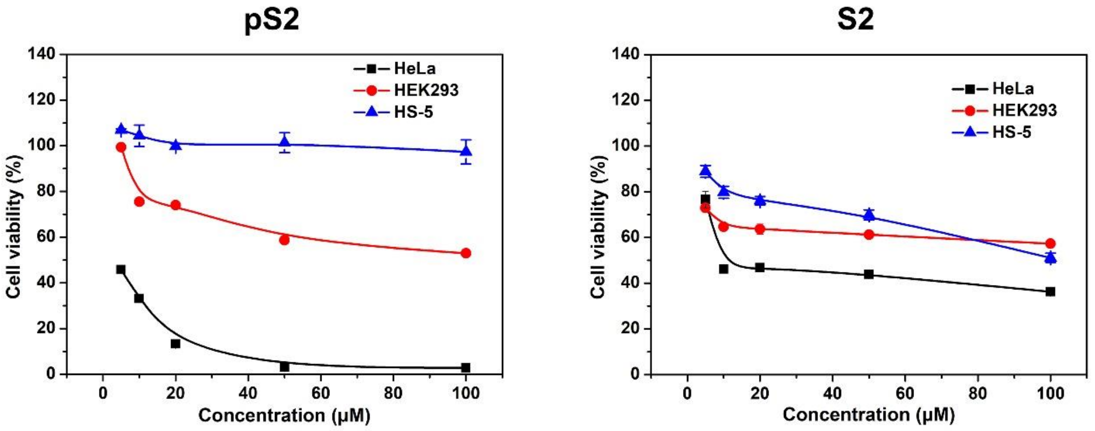


**Figure S19.** Cytotoxicity of **pS2** and **S2** against HeLa cells, HEK293 cells and HS-5 cells for 24 h.

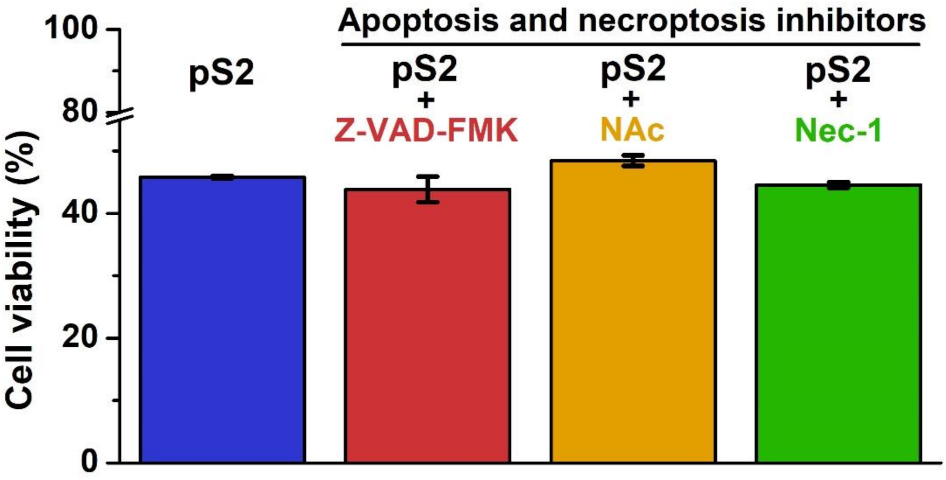


**Figure S20.** Cell viability of HeLa cells treated with only **pS2** (10 μM), the mixture of **pS2** (10 μM) with apoptosis and necroptosis inhibitors (Z-VAD-FMK, 50 μM; NAc, 1mM; Nec-1, 50 μM) for 24 h.

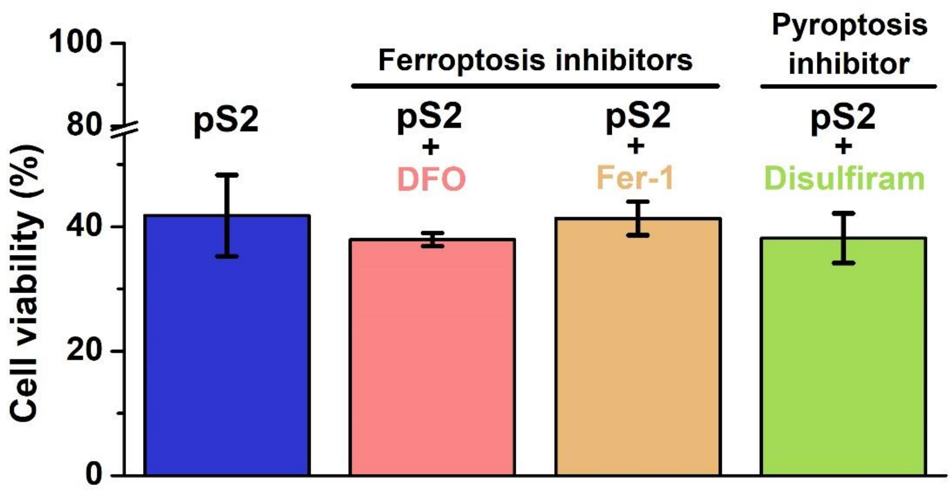


**Figure S21.** Cell viability of HeLa cells treated with only **pS2** (10 μM), the mixture of **pS2** (10 μM) with ferroptosis inhibitors (DFO, 5 μM; Fer-1, 10 μM) and pyroptosis inhibitor (Disulfiram, 5 μM) for 24 h.

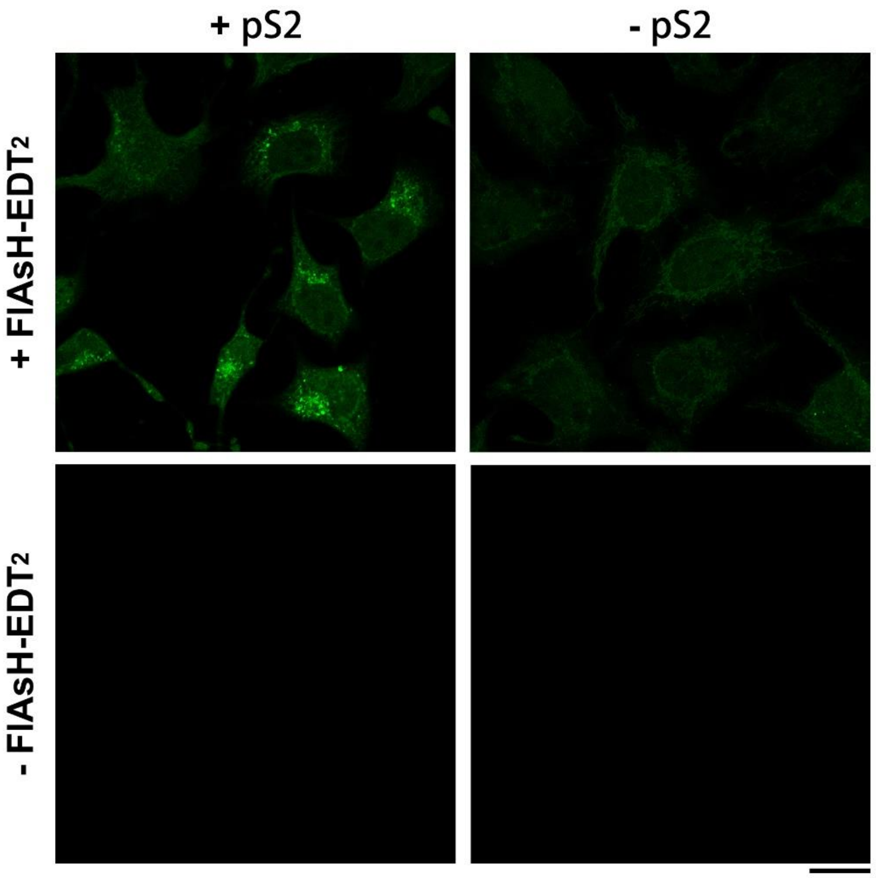

**Figure S22.** CLSM of HeLa cells with or without the pretreatment of **pS2** (10 μM, 30 min) and then with or without staining by FlAsH-EDT2 (1 μM, 1 h). Scale bar = 20 μm.

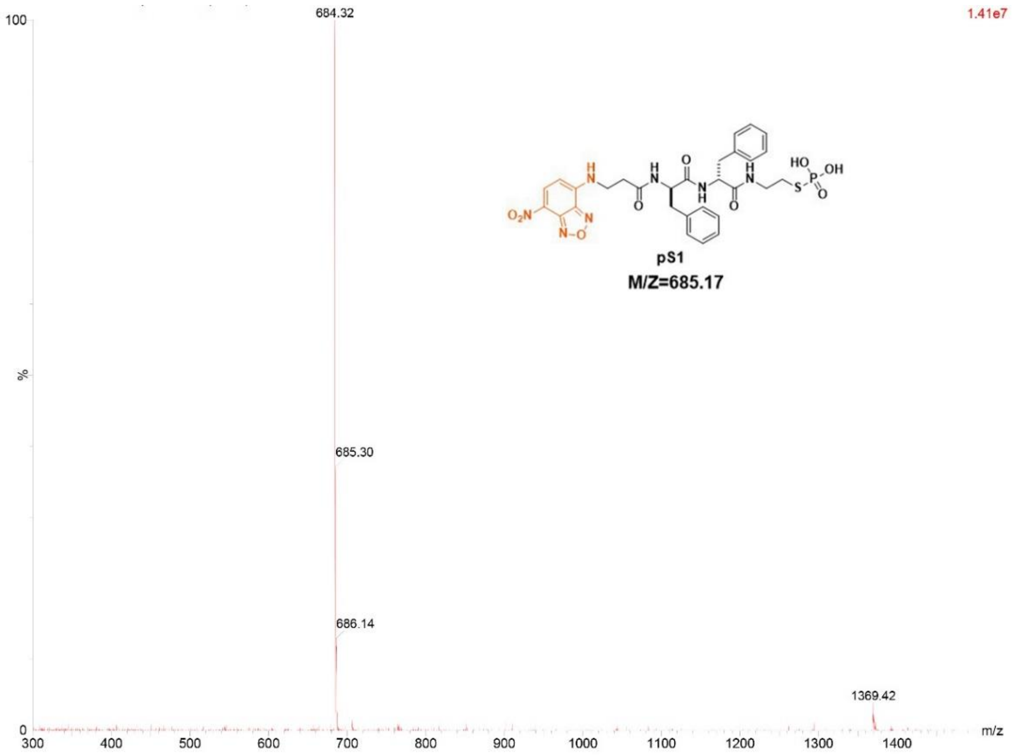


**Figure S23.** LC/MS data for **pS1**. The calculated molecular weight (Mw) of **pS1** is 685.17. The observed M/Z =684.32.

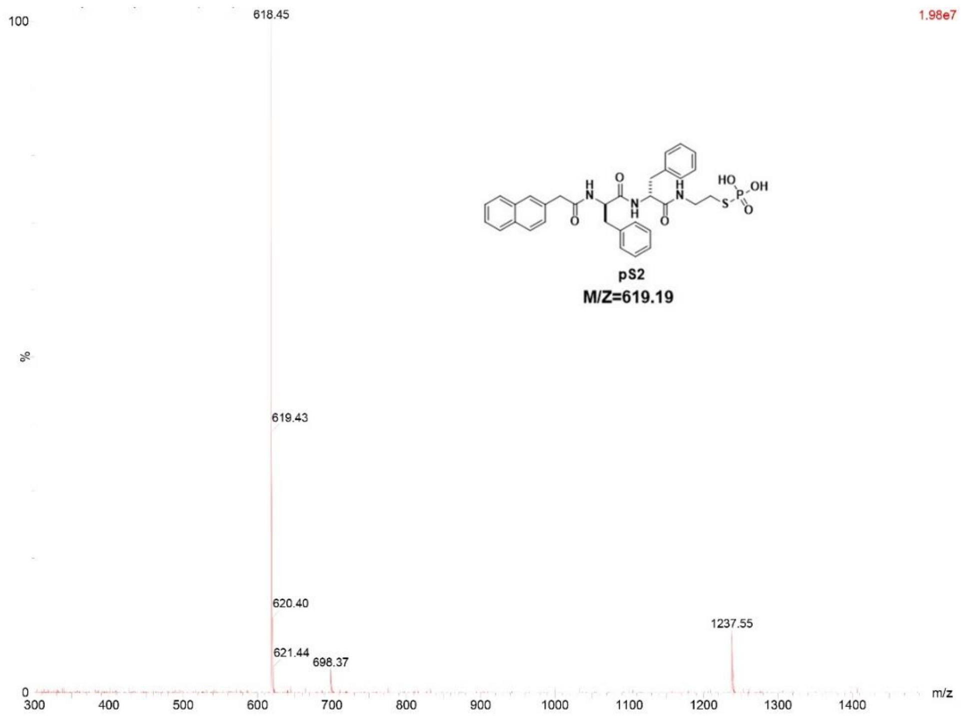


**Figure S24.** LC/MS data for **pS2**. The calculated molecular weight (Mw) of **pS2** is 619.19. The observed M/Z =618.45.

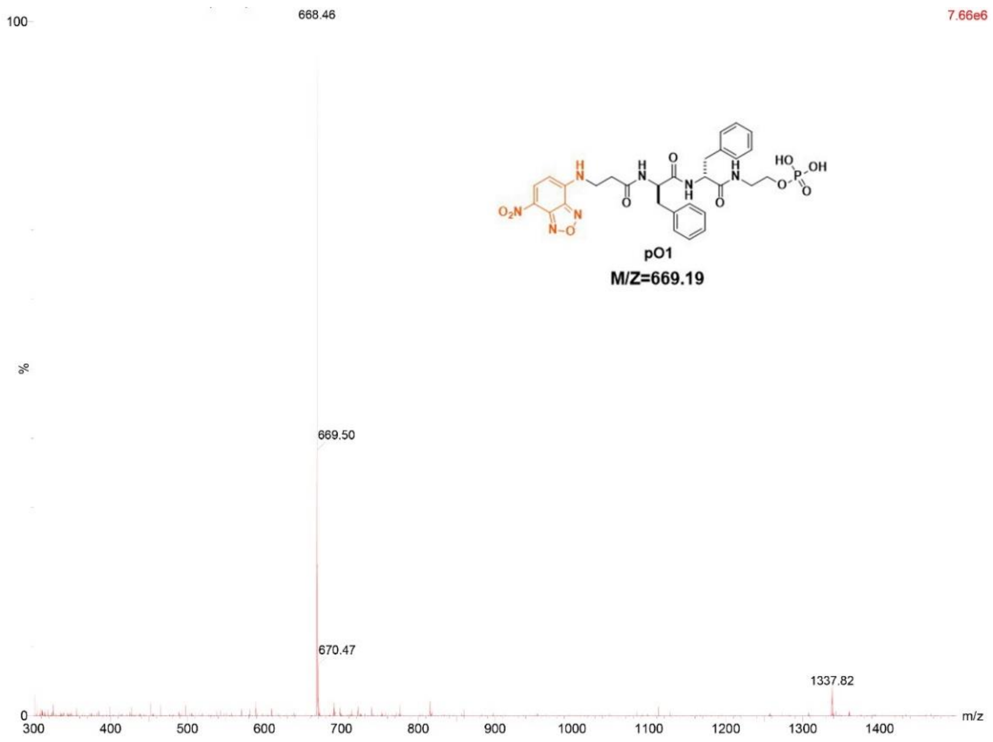


**Figure S25.** LC/MS data for **pO1**. The calculated molecular weight (Mw) of **pO1** is 669.19. The observed M/Z =668.46.

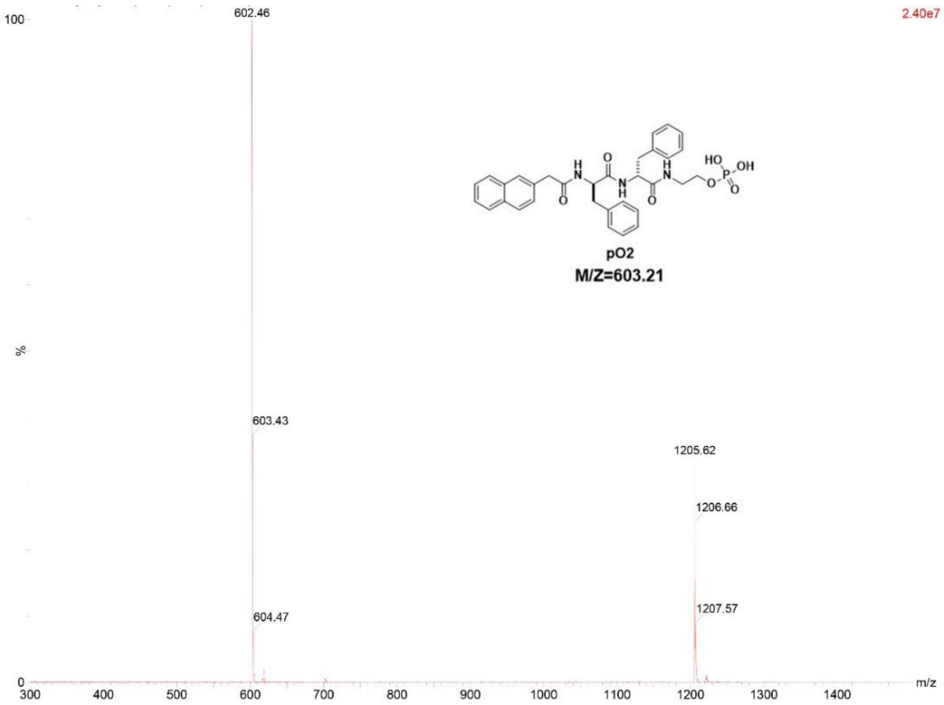


**Figure S26.** LC/MS data for **pO2**. The calculated molecular weight (Mw) of **pO2** is 603.21. The observed M/Z =602.46.
